# INTEGRATED GLOBAL CHICKEN REFERENCE PANEL FROM 13,187 CHICKEN GENOMES

**DOI:** 10.1101/2023.12.12.571301

**Authors:** Di Zhu, Yuzhan Wang, Hao Qu, Chugang Feng, Hui Zhang, Zheya Sheng, Yuliang Jiang, Qinghua Nie, Suqiao Chu, Dingming Shu, Dexiang Zhang, Lingzhao Fang, Yiqiang Zhao, Yuzhe Wang, Xiaoxiang Hu

## Abstract

Chickens are a crucial source of protein for humans and a popular model animal for bird research. Despite the emergence of imputation as a reliable genotyping strategy for large populations, the lack of a high-quality chicken reference panel has hindered progress in chicken genome research. To address this issue, here we introduce the first phase of the 100 K Global Chicken Reference Panel Project (100 K GCRPP). The project includes 13,187 samples and provides services for varied applications on its website (http://farmrefpanel.com/GCRP/). Currently, two panels are available: a Comprehensive Mix Panel (CMP) for domestication diversity research and a Commercial Breed Panel (CBP) for breeding broilers specifically. Evaluation of genotype imputation quality showed that CMP had the highest imputation accuracy compared to imputation using existing chicken panel in animal SNPAtlas, whereas CBP performed stably in the imputation of commercial populations. Additionally, we found that genome-wide association studies using GCRP-imputed data, whether on simulated or real phenotypes, exhibited greater statistical power. In conclusion, our study indicates that the GCRP effectively fills the gap in high-quality reference panels for chickens, providing an effective imputation platform for future genetic and breeding research.

## INTRODUCTION

Chickens are the most abundant bird in the world and the most common source of meat for humans. They were domesticated approximately 8,000 years ago from red jungle fowl (*Gallus gallus*), endemic to tropical South and Southeast Asia (1). The spread of domesticated chickens from Asia to the rest of the world coincided with the establishment of trade routes (2). Currently, the subset of domesticated chickens for human consumption (known as broilers) are so prevalent that they represent how human diets have altered the biosphere. For example, in the early 1950s, the Chicken-of-Tomorrow Program was implemented, significantly increasing broiler growth rates and causing up to a five-fold increase in individual biomass (3). Chicken thus have the potential to be a biostratigraphic marker species of the Anthropocene (4).

The red jungle fowl was the first bird and among the first vertebrates to have its genome sequenced (5). The public catalogue of variant sites (dbSNP version = 106) contains approximately 23.43 million single nucleotide polymorphisms (SNPs) and 2.40 million short insertions and deletions (indels). These resources have greatly contributed to gene discovery in genome-wide association studies (GWAS) (6). Furthermore, thousands of breeding individuals have now been successfully sequenced through advancements in low-coverage whole-genome sequencing (LCS) strategies (7–9), allowing for the identification of gaps in SNP arrays that only include fewer SNPs and facilitating the feature optimization of genomic prediction (GP) (10–12).

A high-quality reference panel enhances the accuracy of genotype imputation across populations and is thus essential for genomics research. Through the 1000 Bull Genomes Project, the cattle industry has published thousands of sequencing datasets that have been applied to GWAS and genome prediction (13). A similar panel would likewise be extremely useful for furthering genetic studies on domestic chickens. Currently, the most comprehensive panel is the Chicken2K project (http://chicken.ynau.edu.cn/index/about/index.html), but this focuses only on genomic diversity across wild jungle fowl and domestic populations, without addressing breeding of commercial broilers. Furthermore, although several multispecies databases include chickens (14,15), the limited amount of species-specific reference haplotypes has hindered accurate genome research in the world’s most popular poultry. Hence, we formulated the 100 K Global Chicken Reference Panel Project (100 K GCRPP). This endeavor aims to discover, genotype, and provide accurate haplotype data for domestic chickens worldwide, including commercial broiler breeds.

In this study, we present results from the first phase of our project. Here, our aim was to compare different strategies for reference panel construction, as well as their applications in genotype imputation and causal gene fine-mapping studies. We conducted low-coverage sequencing of 11,340 broiler individuals (Commercial Breed Panel, CBP), and deep sequencing of 1,847 domestic individuals (Comprehensive Mix Panel, CMP). Our findings offer insight on chicken genetic variation that is more comprehensive and consistent than those of previous studies. The GCRPP will enable us to better understand the landscape of functional variation, genetic associations, and artificial selection in chickens.

## MATERIAL AND METHODS

### Sample Collection and Sequencing

The CBP consisted of 11,340 individuals from five populations, including four yellow-feather broiler lines (YB1, YB2, YB3, YB4) and one white-feather broiler line (WB1). Sample sizes were 3,572, 959, 2,000, 2,809, and 2,000, respectively. All subjects originated from commercial breeding companies and were subjected to low-coverage sequencing at depths ranging from 0.4 – 1.7×. Genomic DNA was extracted from blood samples of every chicken using a DNeasy Blood & Tissue Kit (Qiagen 69506). Extracted DNA quality was assessed in a NanoDrop spectrophotometer and verified with 1% agarose gel electrophoresis. Samples were quantified using a Qubit 2.0 Fluorimeter, then diluted to 40 ng/mL in 96-well plates. A Tn5-based library generation method was employed to produce sequence libraries, as described previously (7). Final libraries were sequenced on the MGISEQ-2000 platform, yielding 2×100 bp paired-end reads.

The CMP used data downloaded from the Sequence Read Archive (SRA), including 1,847 re-sequenced samples across 114 breeds/lines. The samples encompassed both commercial broiler and layer chicken populations, as well as local breeds worldwide (see Supplementary Table 1).

### Genotyping and Haplotype Construction

Raw sequencing reads for low-coverage samples were trimmed using Trimmomatic version 0.36 (16) and subsequently mapped to the GRCg6a (NCBI RefSeq assembly: GCF_000002315.6) reference genome using the GTX-One platform (17). GTX-One is an FPGA-based hardware accelerator optimized for BWA (18) and GATK (19) best-practice workflows, including marking PCR duplicates and indexing BAM files. Quality checks were performed using indel realignment and base quality score recalibration (BQSR) modules within GATK. Next, SNP variants and allele frequency were estimated in BaseVar version 1.01 (20). After filtering out SNPs with a population depth of <1.5-fold interquartile range (IQR) and more than two allele types, 42.23 M candidate SNPs were left for imputation. GeneTalks (China) conducted SNP imputation, using accelerated STITCH (21) software. Parameter K (number of ancestral haplotypes) was set to 20 because five populations were included in the analysis. Post-imputation, SNPs with a minor allele count (MAC) < 5 and an INFO score < 0.40 were excluded, retaining 20.41 M variants.

For the CMP, we downloaded SRA files for each individual and converted them to FASTQ files using SRAtoolkit version 3.0.2. Similar to the CBP, all FASTQ files were processed to generate gVCFs based on GRCg6a, and joint calling was executed for all CMP individuals using the GTX-One platform. SNPs were then filtered using GATK version 4.1.9.0 VariantFiltration with the parameters set to ‘QD < 2.0, MQ < 40.0, FS > 60.0, SOR > 3.0, MQRankSum < −12.5, ReadPosRankSum < −8.0, QUAL < 30’. Indels were filtered using parameters ‘QD < 2.0, FS > 200.0, ReadPosRankSum < −20.0, QUAL < 30’. After quality control, 39.33 M SNPs and 4.79 M indels remained.

Both CBP and CMP variants were phased using Beagle 5.2 (22) to generate haplotypes for all samples. The polymorphic spectrum and potential effects of each SNP are accessible via the “SNP Search” function on the Global Chicken Reference Panel (GCRP) website. Reference panels can also be obtained from the “Download” page for localized analysis.

### Population Genetic Analyses

Population genetic analyses were performed to understand the genetic structure of breeds, as well as provide data on CMP and CBP genetic backgrounds for user genotype imputation. First, phylogenetic relationships of CMP samples were evaluated, and principal component analysis (PCA) was performed to determine any clustering. Samples (n = 1,847) were divided into 12 sub-populations based on origin. The IBS distance matrix within CMP samples was calculated using Plink version 1.9 (23) (--genome; --distance-matrix). This matrix was the foundation for constructing an unrooted neighbor-joining tree in FastME (24). Finally, PCA for the CMP was executed using SMARTPCA (25) based on LD-pruned data derived from Plink (--indep-pairwise 50 10 0.2).

The CBP contained populations with a larger sample size and met broader criteria for genetic analysis. Variant loci and haplotypes shared among the five groups were investigated. To lower the possible impact of genotyping errors on each population, only SNPs with MAC > 3 were considered polymorphic variants in each group. An analysis of SNP, haplotype sharing, and polymorphisms was conducted across populations. Samples in the CBP were then combined with three CMP populations that had sample sizes > 100, including RJF, Indian natives, and Tibet. To quantify nucleic acid diversity, “pi” (26) was computed using vcftools version 0.1.16 (27) (-- pi-window 300k --step 100k) with SNPs shared between CBP and CMP.

For haplotype diversity analysis, overlapping SNP density was further reduced via selecting one SNP from every 20-SNP window and ensuring that SNPs chosen from neighboring windows were approximately equidistant. The analysis utilized 428k SNPs, and a sliding window-based approach was employed to construct haplotypes, Window size was set to eight SNPs, with no overlap between consecutive windows. For each block, haplotype diversity (H) per population was calculated, as described previously (28):

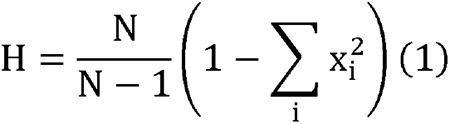

where N is the sample size and xi is the frequency of haplotype i within populations.

### Imputation Assessment

Assessment of imputation effectiveness used 205 CMP samples with >20× sequencing depth as test data (Supplementary Table 3). We used their genotypes obtained from GATK best practice as the gold standard, and later excluded these samples from the CMP for further evaluation. To assess whether GCRP had an advantage in genotype imputation over other panels, chicken panels were downloaded from animal SNPAtlas (14) database as a control.

Considering real imputation practical, two scenarios were tested. The first aimed to assess imputation from low- or high-density arrays to whole-genome sequencing (WGS). Gold-standard GATK results were down-sampled using Affymetrix 600k and Illumina 60k SNP array datasets. Subsequently, imputation was performed using SNPAtlas, CBP, and CMP panels across Beagle5.2, Impute5 version 1.1.5 (29), and Minimac3 version 2.0.1 (30). The second scenario was designed for low-coverage sequencing. Sequencing data at 1× depth were randomly extracted from the original Fastq files for each test sample and subjected to imputation using Glimpse version 1.1.1 (31) and Quilt version 1.0.3 (32).

Accuracy was assessed via computing genotype consistency (GC) for each individual, defined as number of correctly imputed genotypes divided by total number of imputed genotypes. Prior to GC computation, definitive genotypes were assigned to loci based on the highest posterior probability provided by imputation software. Furthermore, allelic dosage (r2) was calculated for every imputed SNP locus (r2 = square of correlation coefficient between imputed allelic dosage and true dosage). All evaluations of genotype imputation accuracy were performed on the medium-sized chromosome 6 to minimize computational complexity.

### Application of GCRP in GWAS

An additional commercial population was used for assessing the utility of GCRP in GWAS. The additional population comprised 4,749 samples from the same generation, all subjected to low-coverage sequencing. Genotype imputation was performed with the CBP and Glimpse. After imputation, loci were filtered using minor allele frequency (MAF) > 0.01 and INFO score > 0.4, leaving 9.5 M SNPs for GWAS.

Both simulated phenotypes and real anonymized phenotype A were used in GWAS. Simulated phenotypes were generated from real genotypes in R package SIMER (https://github.com/xiaolei-lab/SIMER), based only on additive genetic and residual effects. Heritability was set at 0.3, with 100 causal mutations randomly selected from all included SNPs. Their effects were sampled from the default normal distribution of SIMER. Residual terms were sampled from another normal distribution with variance adjusted to heritability.

The mixed linear model used in GWAS was fitted using the GCTA version 1.92.0 “mlma” (33):

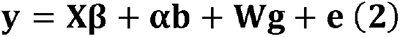

where **y** is a vector of phenotypes; for the real trait β is a vector of fixed effects, including batch and chicken coop effects; for the simulated trait β is only include overall mean; **b** is a vector of genotypes (coded 0, 1, and 2); α is the effect of test SNP; **g** is a vector containing random polygenic effects; 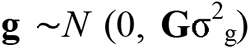; **G** is the GRM constructed using LCS SNPs; 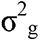 variance explained by SNPs; **e** is a vector of random residual effects and 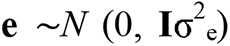; **I** is the identity matrix; **X** and **W** are design matrices connecting phenotypes to fixed effects and random polygenic effects, respectively. **X** is the identity matrix of the simulated trait.

## RESULTS

### Global Chicken Reference Panels

We generated the CMP using 1,847 individuals from 114 breeds spanning various continents (Supplementary Table 1). Additionally, we generated a CBP specifically for chicken breeding, encompassing 11,340 individuals from five major commercial breeds (Supplementary Table 2). The two panels combined formed the pilot phase of the GCRP database (Figure 1).

**Figure 1.**
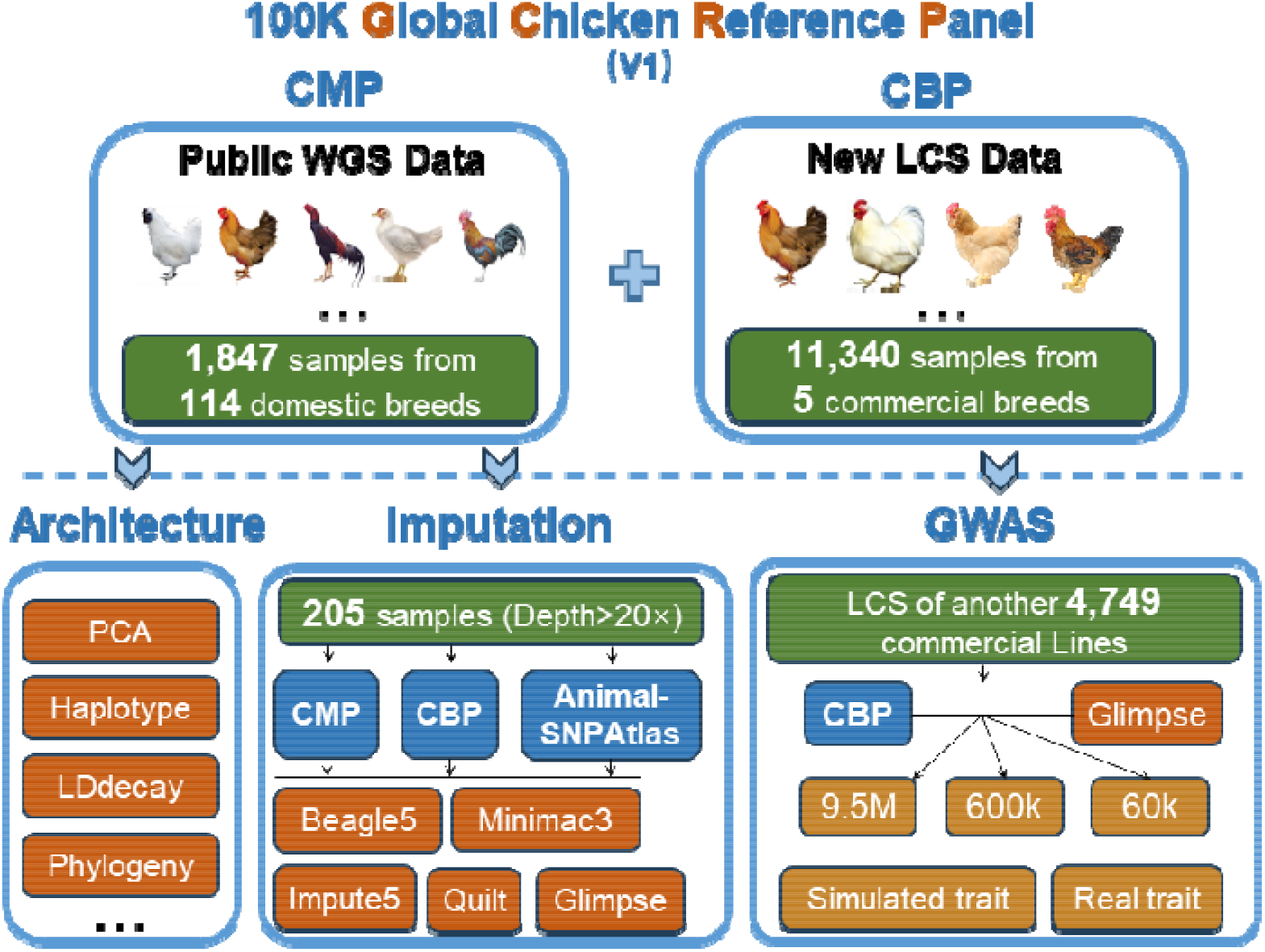
Flow chart of the Global Chicken Reference Panel (GCRP) database and study design. CMP (Comprehensive Mix Panel) is a domestication panel for thousands of chicken breeds and CBP (Commercial Breed Panel) is an improvement panel for tens of billion commercial chickens. WGS, whole genome sequencing; LCS, low-coverage sequencing.

Overall, we identified 48.02 M SNPs and 4.79 M indels in GCRP, including 26.36 M rare SNPs and 3.53 M rare indels (0.0001 < MAF < 0.01). Notably, while 66.40% of SNPs in dbSNP were present in GCRP, we discovered 32.47 M novel SNPs and expanding the chicken dbSNP by 139% (Supplementary Figure 1). Moreover, 42.59% of SNPs (8.69 M) in CBP did not overlap with those in CMP, indicating that significant changes resulted from artificial selection. Only a small portion (5.42%) of indels in CMP overlapped with those in dbSNP (Supplementary Figure 1). It is worth noting that CBP temporarily excluded Indels due to the limitations of low-coverage sequencing.

### Population Structure Analyses

For the CMP, we categorized the 114 populations into 12 origin-based subgroups (Supplementary Table 1). The results of PCA and phylogenetic analyses revealed that most individuals from the same origin tended to cluster together (Figure 2a and b). Overall, the domestication trajectory from red jungle fowl to domestic and commercial chickens was clearly visible. Domestication began in South Asia and spread through pathways that reached Southeast, Northeast, and West Asia, then eventually Europe. Additionally, red jungle fowl continued to differentiate and became more distantly related to the other groups.

**Figure 2.**
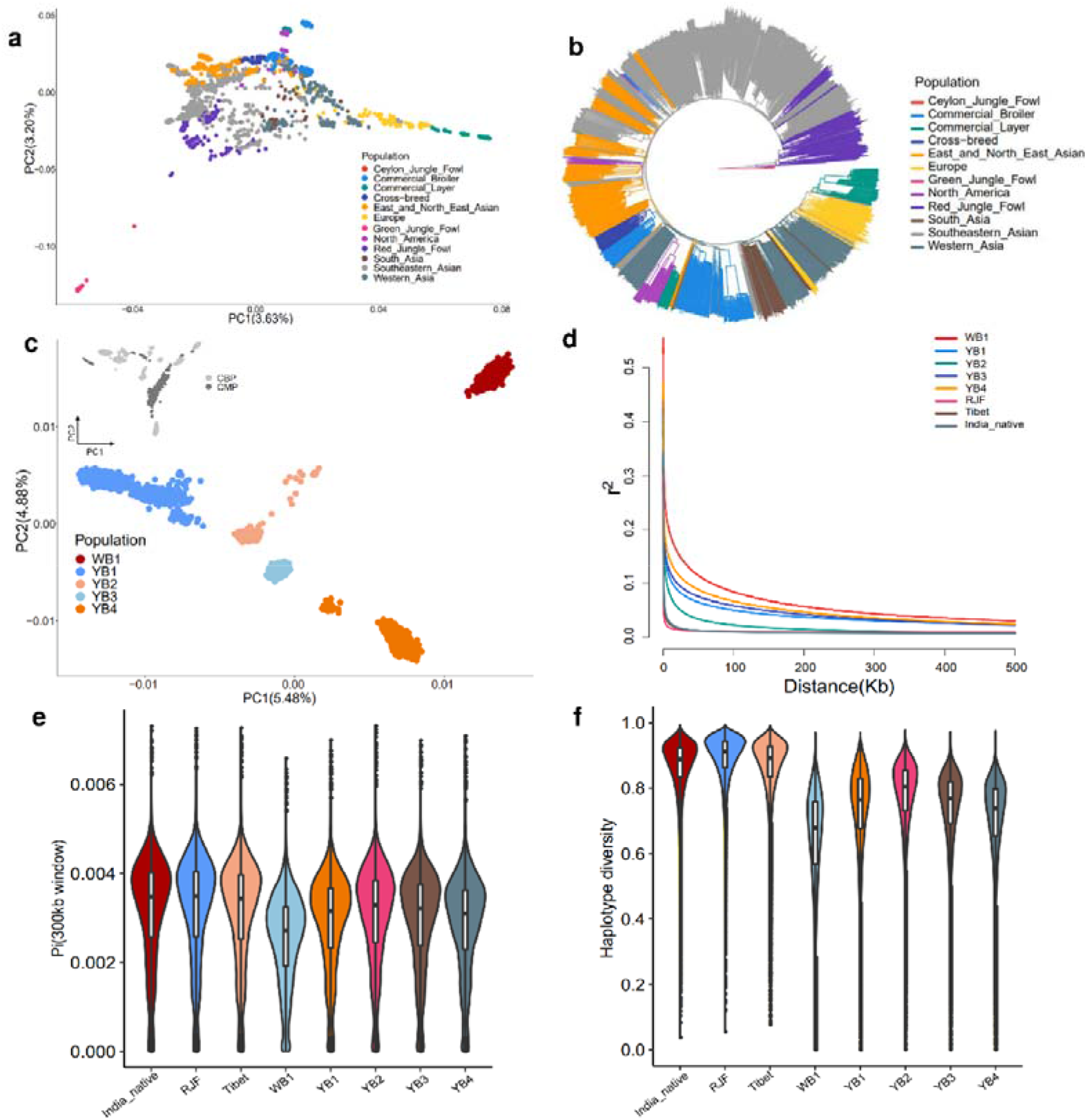
Population genetics of GCRP samples. (a) Principal component analysis (PCA) for CMP, with 12 subgroups represented by different colors. (b) Phylogenetic analysis of CMP, with 12 subgroups indicated by different colors. (c) PCA for CBP; the top-left corner illustrates the relationship between CBP and CMP through overlapping loci in two panels. (d) Linkage disequilibrium decay after merging CBP with the three largest CMP populations. (e) Violin plots displaying nucleotide diversity (π) distribution within 300 kb windows for each population. (f) Violin plots showing haplotype diversity distribution per population.

The CBP samples segregated clearly according to breed and were significantly distinguishable from domesticated breeds in the CMP (Figure 2c). Most SNPs were shared among all five populations, but only 8.89% of haplotypes were shared across all groups (Supplementary Figure 2). This pattern indicates that artificial breeding of specific traits shaped the genomes of different broilers. Further population structure analysis on all CBP populations and three populations in CMP revealed higher nucleotide polymorphism, greater haplotype diversity, and faster LD decay in domestic breeds (Indian native and Tibet) and red jungle fowl than in the commercial breeds (Figure 2d–f). Within the CBP, the YB2 population had the highest variant and haplotype diversity, whereas the WB1 population had the lowest (Figure 2e and f). Haplotype polymorphism was high YB1, while SNP polymorphism was moderate. These differences in genome architecture accurately reflected breeding stages: YB1 was in the hybrid breeding stage, YB2 was in the conservation stage, and WB1 was in the purebred breeding stage.

### Assessment of Genotype Imputation

Our imputation quality assessment showed that sequencing-based strategies had considerably higher genotype consistency (Figure 3a–c) and dosage r^2^ (Supplementary Figure 3a) than two SNP array-based strategies. The highest genotype consistency (99.4%) and dosage r^2^ (0.961) were obtained using the CMP and glimpse software to impute low coverage sequencing data. Among array-based imputation results, Affymetrix 600k performed better than Illumina 60k. When using CMP as reference panel, the GC for three imputation softwares targeting the Affymetrix 600k was found to be 0.924, 0.926, and 0.885, respectively. These values were higher compared to the Illumina 60k, where the GC rates were 0.800, 0.810, and 0.748, respectively (Figure 3a-b). Allele dosage r^2^ for each estimated SNP aligned with the trends for GC (Supplementary Figure 3).

**Figure 3.**
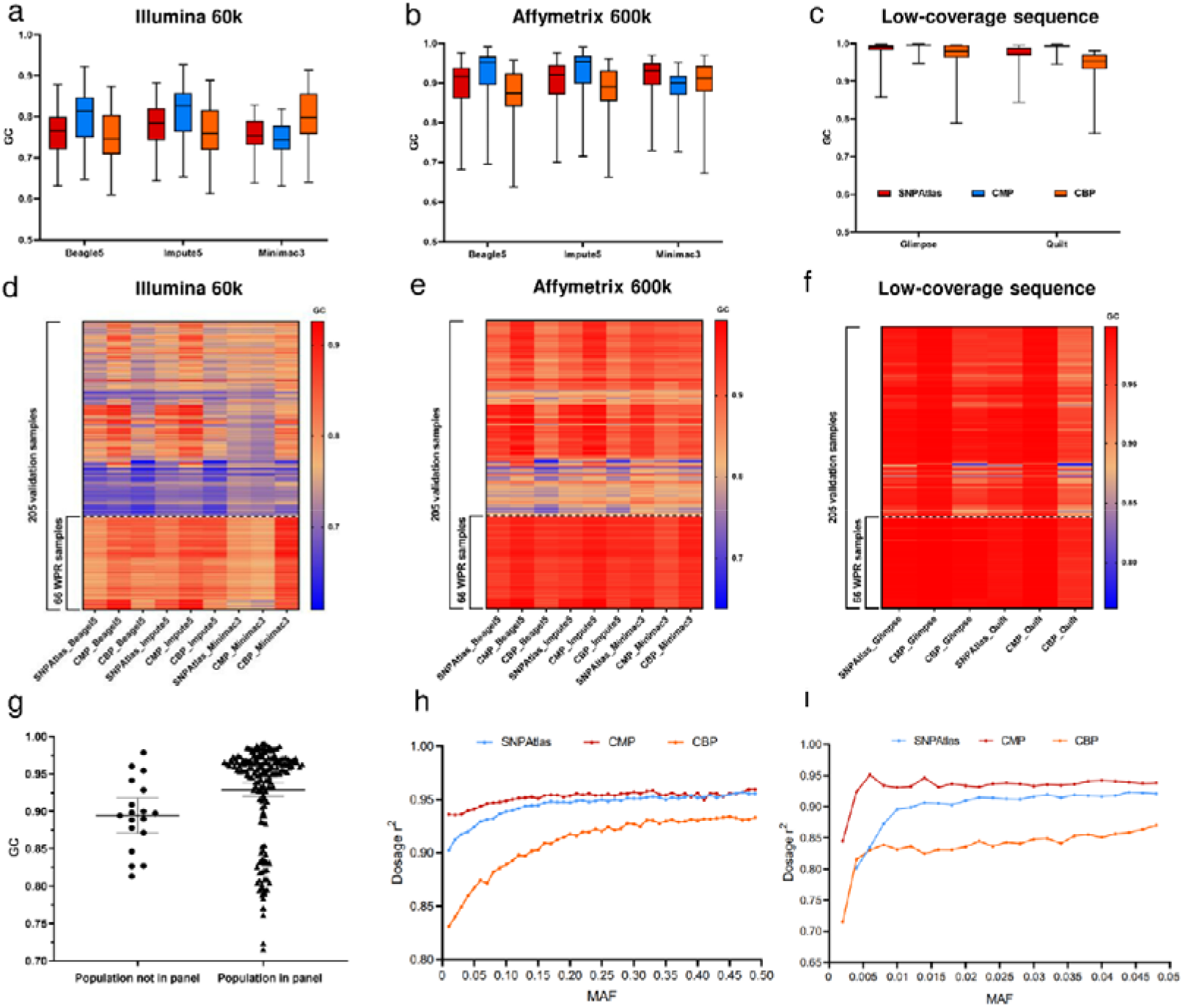
Assessment of genotype imputation accuracy. Boxplots represent genotype concordance of imputed versus observed genotypes. The assessment used three different software and three reference panels: Illumina 60k array (a), Affymetrix 600k (b) and low-coverage sequencing (c). Color coding is consistent across a, b, and c. Heatmaps representing imputed genotype concordance of 205 test samples: Illumina 60k array (d), Affymetrix 600k (e) and low-coverage sequence (f). The 66 commercial white Plymouth rock (WPR) samples are displayed at the bottom of each heatmap. (g) Comparison of genotype imputation accuracy between “population in panel” and “population not in panel” samples (P = 0.004, one-sided Student’s t test). (h, i) Imputation accuracy calculated in Glimpse: (h) minor allele frequency (MAF) bins (0.01 bin size) from 0 to 0.5 and (i) MAF bins (0.002 bin size) from 0 to 0.05.

Comparison of the three panels showed that CMP had the highest accuracy in 26 of 27 imputation strategies, followed by SNPAtlas, and finally by CBP (Figure 3a–c and Supplementary Figure 3a). However, we observed that 66 commercial white Plymouth rock (WPR) population-derived samples out of the 205 samples exhibited higher accuracy in CBP compare with others domestic samples, especially with the low-density 60k array (Figure 3d-f, supplementary figure 3b and supplementary figure 4). This result was consistent with our expectation that CMP and CBP would serve distinct roles in research on domesticated and commercial broiler breeds, respectively. We divided the 205 samples into two groups, one containing individuals from the same population in the remaining CMP samples (population in the panel) and the other containing no individuals from the same population (population not in the panel). The results demonstrate that the accuracy of related individuals is significantly higher than that of unrelated individuals (Figure 3g), highlighting the crucial influence of the relationship between the panel and the target sample on imputation accuracy. Furthermore, SNPs with lower MAF tended to have lower dosage r^2^ (Figure 3h-i, supplementary figure 5). In the evaluation of rare mutations (0.002 < MAF < 0.05), GCRP successfully detected rare mutations exceeding 0.005 (Figure 3i).

### Case Study of GWAS for Chicken Complex Trait

Increasing marker density using imputation based WGS data can enhance the statistical power of GWAS to detect associated signals (34–36). We employed CBP to perform genotype imputation on an external commercial population of 4,749 individuals and obtained a set of 9.5 M SNPs. We then compared three SNP sets (Illumina 60k, Affymetrix 600k, and our imputed 9.5 M set) in GWAS performance using both simulated and real anonymous phenotypes. Simulated phenotypic heritability was set at 0.3, and we identified seven independent causative mutations exceeding the significance threshold (5 × 10^-8^), representing seven segregating quantitative trait loci (QTL). All three SNP sets successfully captured the most significant QTL on chr18 (Figure 4a). The 600k array and 9.3 M set covered all significant QTL (false negative rate, FNR = 0), but the 60k array missed two QTL intervals (FNR = 28.57%). Additionally, the 9.3 M set revealed that the average distance between the top SNP within each QTL and the causative mutation was 1.2 kb (range: 0–5.3 kb), significantly shorter than for the other two sets (60k: 69.1 kb, 600k: 75.3 kb).

**Figure 4.**
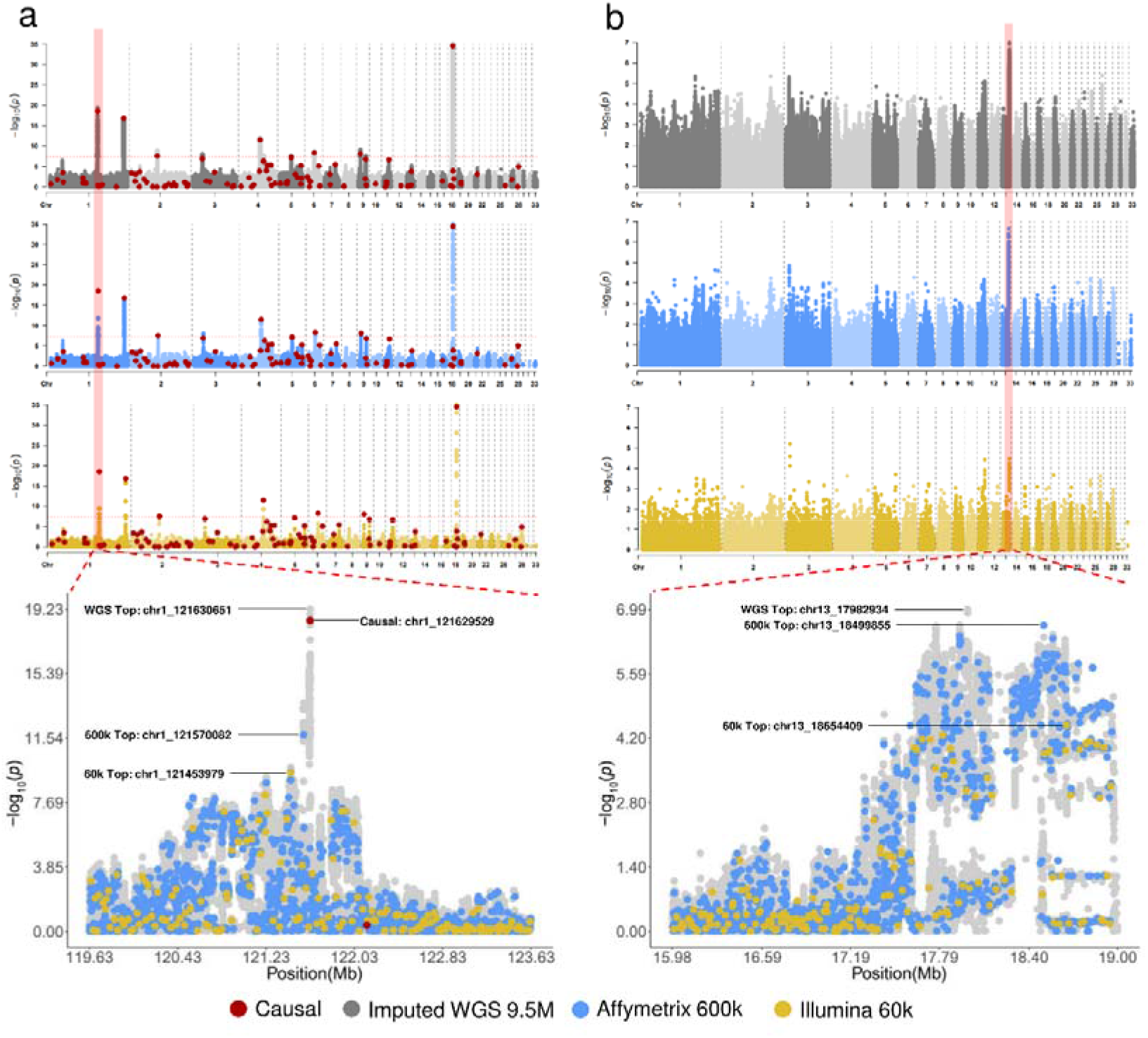
Application of GCRP in genome-wide association studies. Manhattan and zoom plots for simulated trait (a) and real anonymized phenotype (b). The −log10 (P-value) scale (y-axis) are plotted for each SNP position (x-axis). Colors reflect different SNP sets. Red dots indicate locations of true causal mutations in simulated data.

We observed similar findings in the real phenotype (*h^2^*= 0.24), where the 9.3 M set exhibited stronger significance than the two arrays. Only the 9.3 M set and the 600k array detected the largest peak on chr13 (chr13: 17.41–18.97 Mb, Figure 4b). Our results substantiate the hypothesis that GCRP provides distinct advantages in GWAS over low-or high-density SNP arrays.

### GCRP Web-based Services

The Global Chicken Reference Panel offers a user-friendly interface (http://farmrefpanel.com/GCRP/#/), comprising several functional modules (Figure 5). First, ‘variantDB’ allows users to search and visualize detailed data on variants in CBP or CMP, including site position, annotation, and allelic frequency in different populations. Second, the ‘Imputation’ module enables users to run online genotype imputation using three imputation tools: Beagle5, Minimac3, and Impute5. Here, users can upload their VCF and download the imputation results. Finally, the download module allows users to download VCF files for both CBP and CMP. Users can access these modules by clicking ‘Download’ button to download GCRP VCF files and the corresponding buttons on the ‘Home’ page. The ‘About’ page provides a comprehensive description of the panel’s purpose and components, along with a graphical display of sample geographical distribution, evolutionary distance, and relationships. Finally, users can submit feedback through the email addresses provided on the ‘Contact Us’ page.

**Figure 5.**
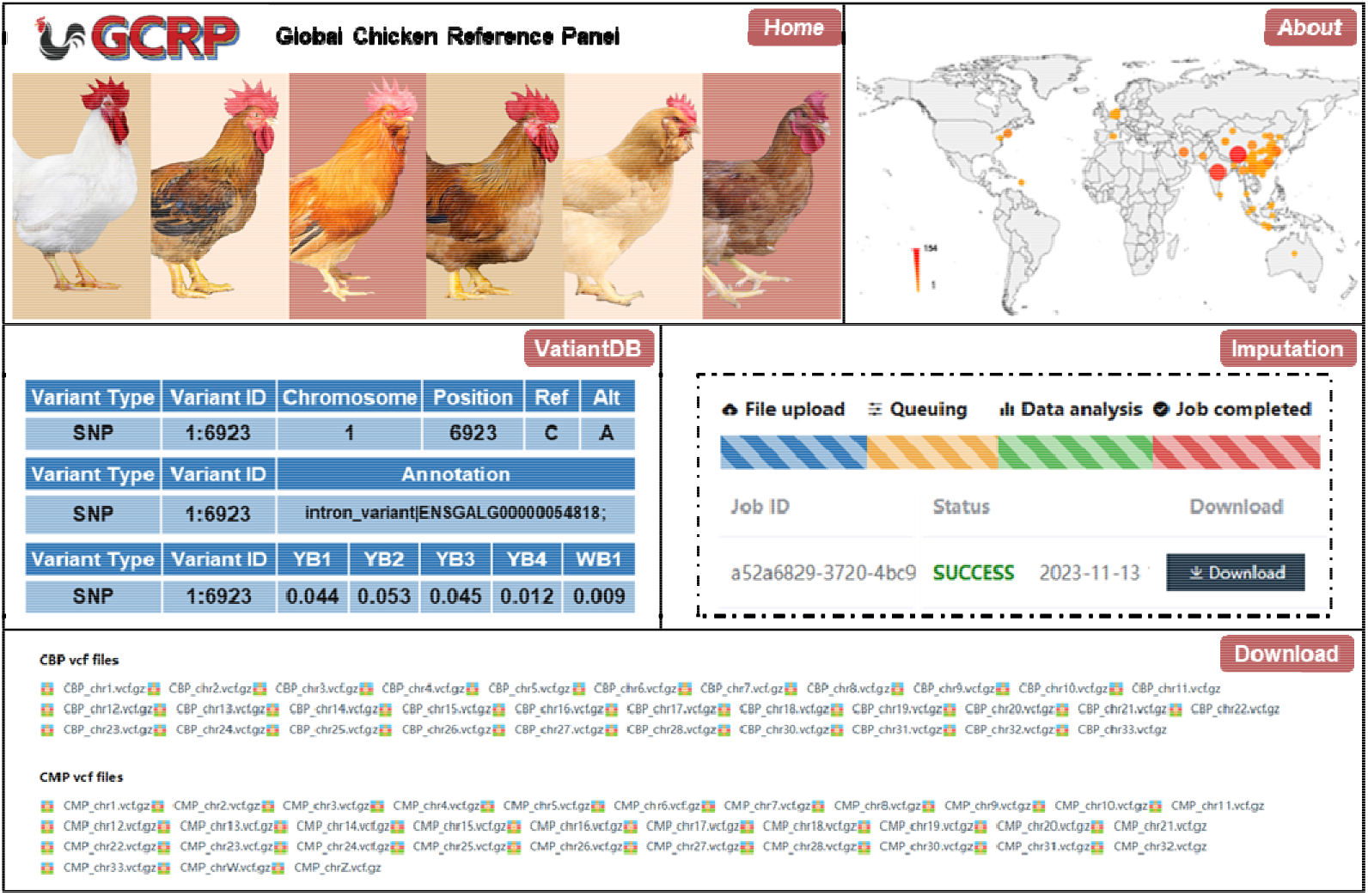
Summary of GCRP Web-based Services. Main modules in GCRP are ‘Home’, ‘About’, ‘VariantDB’, ‘Imputation’, and ‘Download’. The ‘About’ page presents the geographical distribution of breeds collected in this study. The circle sizes and colors correspond to the sample size for each study area, indicating the number of unique breeds. On the ‘VariantDB’ page, various types of specific information about the recorded variants are displayed, such as allele coordinates, gene annotations, and minor allele frequencies across different breeds. The ‘Imputation’ page showcases the different stages and states of online imputation.

## Discussion

To the best of our knowledge, this is the first study to release a dedicated chicken reference panel and genotype imputation web service. In consideration of different user needs, we launched CBP for precise breeding of broilers and CMP for broader scientific research, making GCRP the most comprehensive chicken genotype database released to date.

The GCRP identified a large number of new variants, including 56% rare mutations (MAF ≤ 0.01), enriching the current chicken dbSNP database. The results of PCA and phylogenetic analyses on CBP and CMP indicated that the samples clustered effectively based on population or origin. Analyses of SNPs, haplotype diversity, and LD decay all highlighted differences in genomic background between commercial and local populations. Furthermore, we discovered that haplotype between populations were more pronounced than those at individual loci, consistent with the historical pattern of relatively isolated evolution among these groups. Overall, our results confirmed the high reliability and quality of GCRP.

Genotype imputation is now the standard in genetic research. Comprehensive human genotype imputation panels and platforms, such as 1000 Genomes Project (37) and NyuWa in China (38), have been provided. For agricultural animals, universal reference panels have been released for cattle (13,39,40), pigs (35,41), and aquatic animals (42). The Animal Impute DB version 2 provides genotype imputation services for 20 animal species, but the sample size of chickens (n=509) was limited and insufficiently representative (14). Compared with the panels in Animal-SNPAtlas, GCRP (n = 13,187) exhibited significant advantages in both array-based and low-coverage sequencing imputation. Results from simulated and real phenotypes showed that post-imputation GWAS yielded stronger signals and higher QTL detection power than two arrays. In summary, our findings provide empirical evidence demonstrating the application potential of GCRP in GWAS.

The CBP resulted in lower imputation accuracy for domesticated chicken breeds than CMP and SNPAtlas; however, CBP performed well when inferring commercial breeds, particularly in comparison with low-density arrays. Using CBP to impute the progeny of included commercial lines achieved 99.9% GC. This finding aligns with our other results (Figure 3d–f) and again illustrates the improvement in accuracy when using large-scale panels of the target population for imputation. The availability of our 100 K GCRPP now makes this strategy less challenging than before. Our results also indicate that the imputation strategy based on low-coverage sequencing outperforms those based on low/high-density arrays, suggesting a more promising application perspective for low-coverage sequencing. However, it is important to note that CBP constructed based on low-coverage sequencing cannot capture Indels currently. Considering the cost-sensitivity of breeding applications, low-coverage sequencing + imputation may still be the most cost-effective research solution for the near future.

In conclusion, our goal was to develop GCRP as a benchmark for future research on chicken population genetics and demonstrate workable methods. This first phase of the GCRP covered hundreds of breeds across multiple continents. As more data accumulate, both CBP and CMP will be continuously updated. For instance, African local breeds and commercial laying breeds will be included to provide a comprehensive panel for chicken genotype imputation in GCRP version 2. This extensive collection will serve as a valuable genetic database for studying the evolution and improvement of chickens and other birds. Additionally, our project will collaborate with the FAANG project (43) and gradually integrate the GCRP with regulatory elements atlas (44), and ChickenGTEx (45). This integration can then be applied to meta- and colocalization analyses (46,47), facilitating a more comprehensive analysis of gene regulatory mechanisms. Multi-omics integration will also contribute to the development of accurate artificial intelligence algorithms (48,49).

## DATA AVAILABILITY

Both CBP and CMP is freely available to the public without registration or login requirements (http://farmrefpanel.com/GCRP/#/.

## AUTHOR CONTRIBUTIONS

Yuzhe Wang: Conceptualization, Project administration, Supervision, Writing-review & editing, Methodology, Funding acquisition. Di Zhu: Writing-original draft, Data curation, Formal analysis, Methodology, Visualization, Writing–review & editing, Software. Yuzhan Wang: Formal analysis, Visualization. Hao Qu: Data curation, Resources. Chungang Feng: Writing-review & editing, Methodology. Hui Zhang: Writing–review & editing, Data curation. Zheya Sheng: Data curation. Yuliang Jiang: Writing–review & editing. Qinghua Nie: Writing–review & editing. Suqiao Chu: Data curation, Resources. Dingming Shu: Writing–review & editing, Resources. Dexiang Zhang: Data curation, Resources. Lingzhao Fang: Supervision, Writing-review & editing. Yiqiang Zhao: Supervision, Writing-review & editing, Data curation. Xiaoxiang Hu: Funding acquisition, Supervision, Writing-review & editing.

## ACKNOWLEDGEMENTS

The authors thank Chouxian Ma, Long Fang and Libing Sun for providing the services of hardware accelerator platform. The authors also thank Jiangli Ren, Yini Jia, Linlin Zhang, Ce Wang, Jiatong Yu, Ran Song for their genomic sequencing and data analyses. The analysis was performed on the high-performance computing platform of the State Key Laboratory of Agrobiotechnology and the National Research Facility for Phenotypic and Genotypic Analysis of Model Animals.

