## supplemental figures for "INTEGRATED GLOBAL CHICKEN REFERENCE PANEL FROM 13,187 CHICKEN GENOMES"

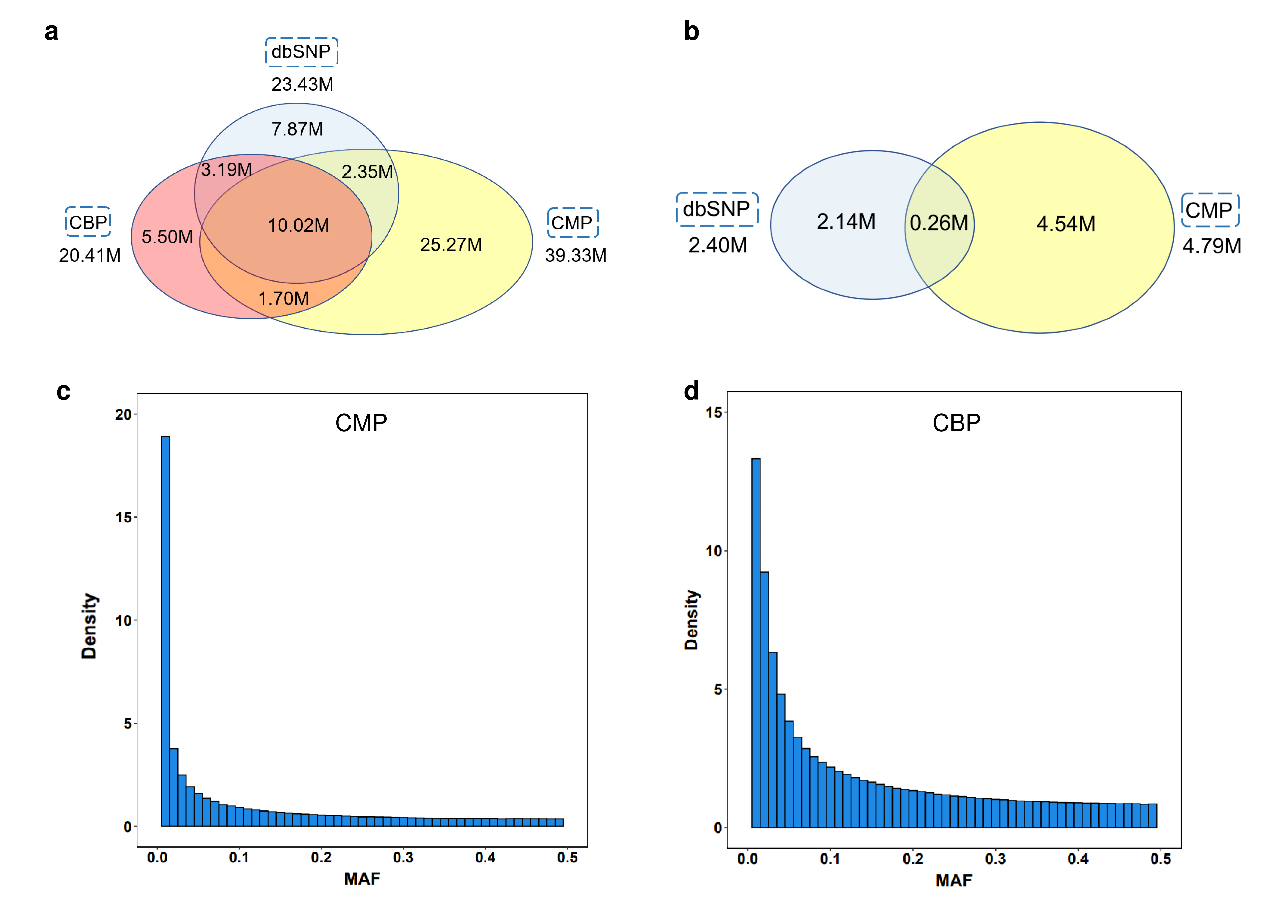

Supplementary figure 1 (a) Venn diagram depicting the overlap of SNP among dbSNP, CBP, and CMP. (b) Venn diagram depicting the overlap of Indel among dbSNP and CMP. (c) Histogram showing the distribution of variant minor allele frequencies in CMP.

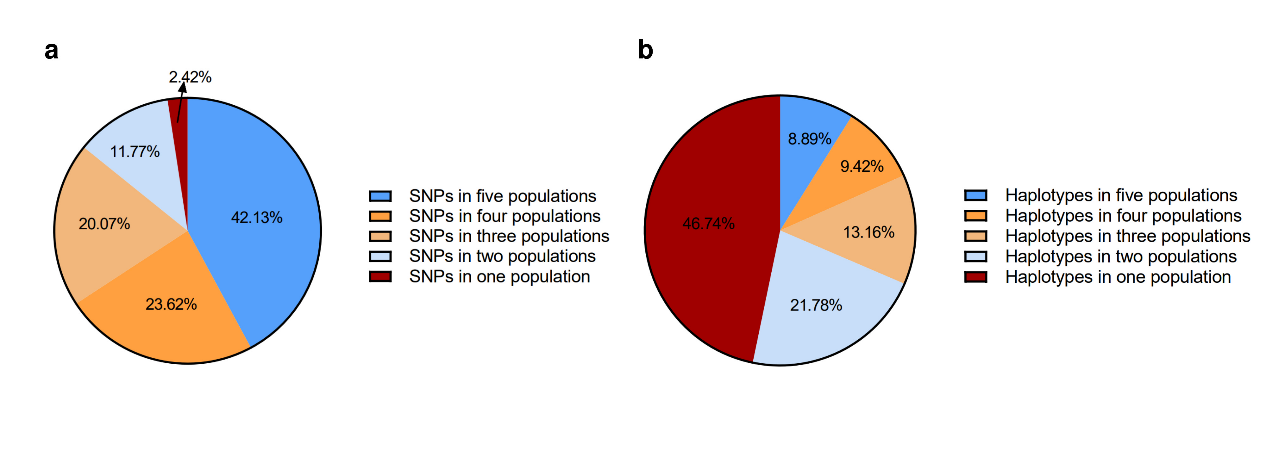

Supplementary figure 2 (a) Proportion of shared SNPs in CBP across five populations. (b) Proportion of shared haplotypes in CBP across five populations.

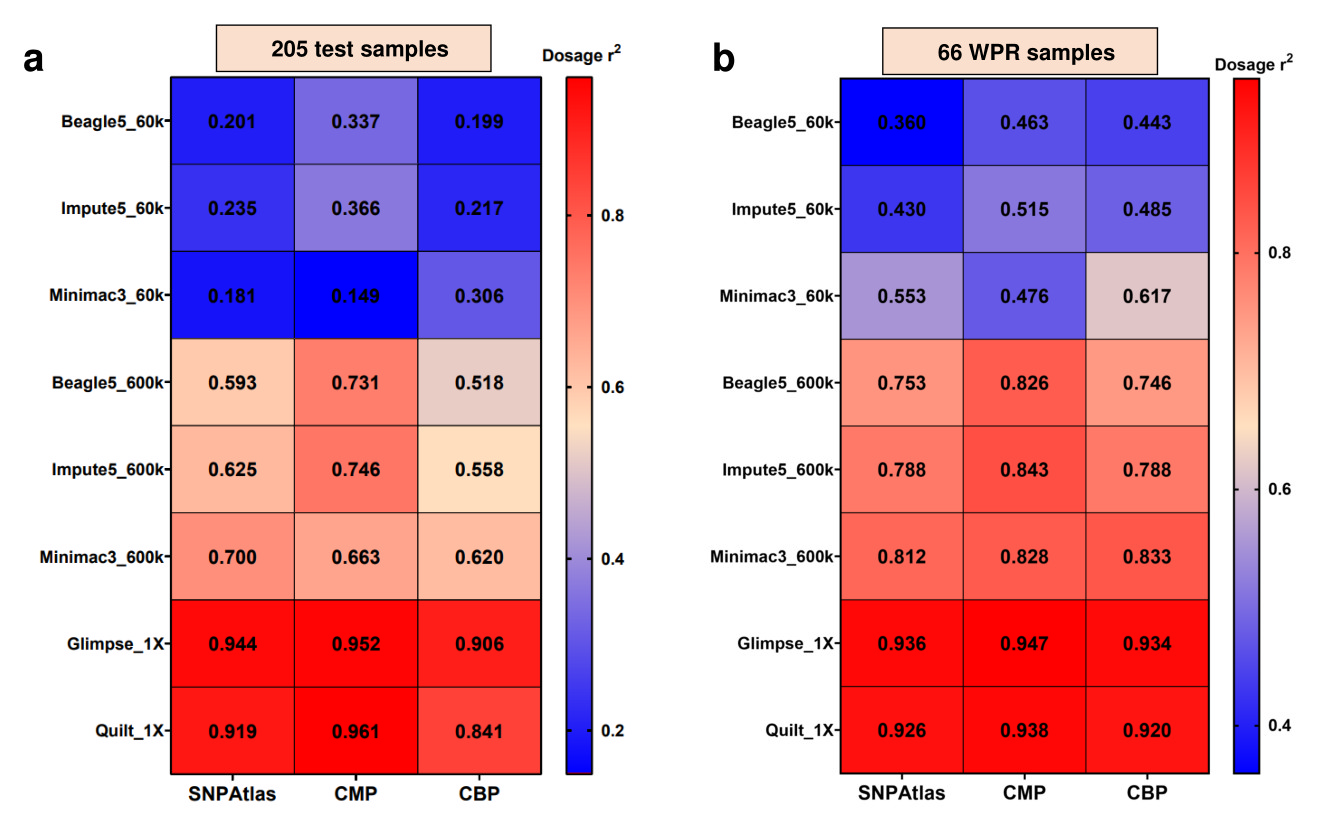

Supplementary figure 3 Square of the correlation between the imputed allele dosage and the true allele dosage for each panel and imputation strategy combination for 205 test samples (a) and 66 commercial white Plymouth rock (WPR) samples.

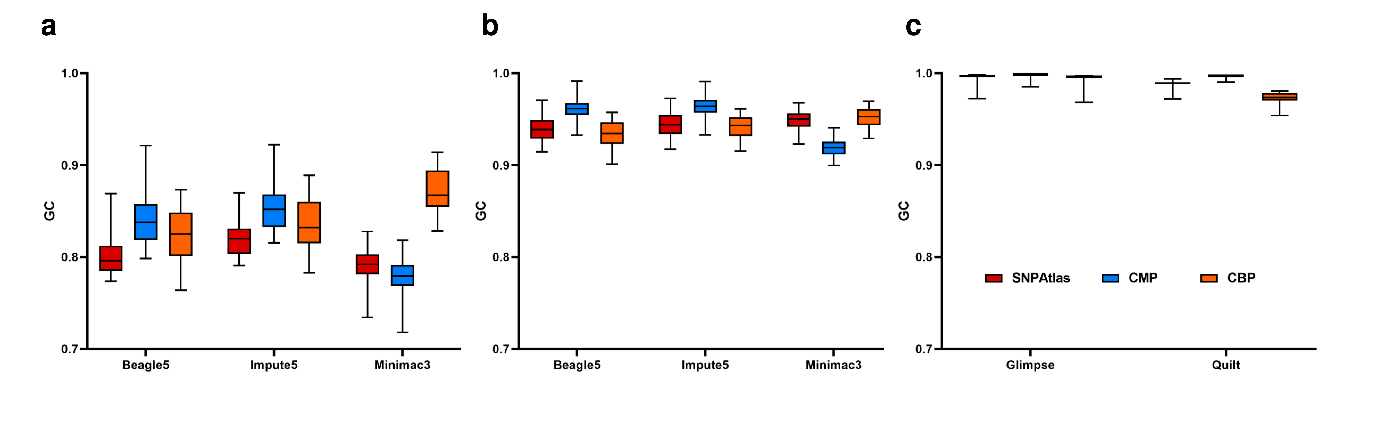

Supplementary figure 4 The genotype concordance of imputed versus observed genotypes using different software and three reference panels targeted to Illumina 60k array (a), Affymetrix 600k (b) and low-coverage sequence (c) for 66 commercial white Plymouth rock (WPR) samples.

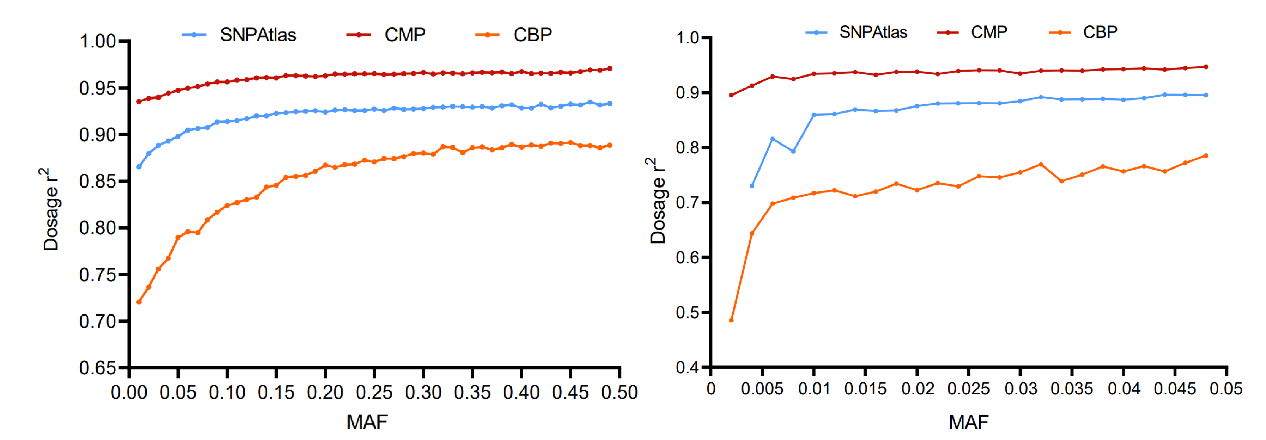

Supplementary figure 5 The imputation accuracy with different MAF bins using Quilt.
